# Single-cell multimodal profiling of monocytes reveals diverse phenotypes and alterations linked to cardiovascular disease risks

**DOI:** 10.1101/2024.02.18.580913

**Authors:** Alexander C. Bashore, Chenyi Xue, Eunyoung Kim, Hanying Yan, Lucie Y. Zhu, Huize Pan, Michael Kissner, Leila S. Ross, Hanrui Zhang, Mingyao Li, Muredach P. Reilly

## Abstract

Monocytes are a critical innate immune system cell type that serves homeostatic and immunoregulatory functions. The Cell surface expression of CD14 and CD16 has historically identified them, however, recent single-cell studies have uncovered that they are much more heterogeneous than previously realized. We utilized cellular indexing of transcriptomes and epitopes by sequencing (CITE-seq) and single-cell RNA sequencing (scRNA-seq) to describe the comprehensive transcriptional and phenotypic landscape of 437,126 monocytes. This high-dimensional multimodal approach identified vast phenotypic diversity and functionally distinct subsets, including IFN-responsive, MHCII^hi^, monocyte-platelet aggregates, and non-classical, as well as several subpopulations of classical monocytes. Using flow cytometry, we validated the existence of MHCII^+^CD275^+^ MHCII^hi^, CD42b^+^ monocyte-platelet aggregates, CD16^+^CD99^-^ non-classical monocytes, and CD99^+^ classical monocytes. Each subpopulation exhibited unique functions, developmental trajectories, transcriptional regulation, and tissue distribution. Moreover, we revealed alterations associated with cardiovascular disease (CVD) risk factors, including race, smoking, and hyperlipidemia, and the effect of hyperlipidemia was recapitulated in mouse models of elevated cholesterol. This integrative and cross-species comparative analysis provides a unique resource to compare alterations in monocytes in pathological conditions and offers insights into monocyte-driven mechanisms in CVD and the potential for targeted therapies.

**Summary:** Multimodal profiling provides a comprehensive phenotypic and transcriptional understanding of monocytes in health and cardiovascular disease risk states.

## Introduction

Blood monocytes, a pivotal component of the innate immune system, play a critical role in immune surveillance, host defense, and inflammation regulation. These versatile cells serve as sentinels, patrolling the circulatory system and differentiating into macrophages and dendritic cells upon reaching inflamed tissues *(1)*. While monocytes have long been regarded as a relatively homogenous population, it has become increasingly evident that they exhibit a substantial degree of heterogeneity in terms of surface marker expression, functional properties, and responsiveness to microenvironmental cues *(2–4)*. This intricate landscape of monocyte subpopulations has significant implications for our understanding of both physiological and pathological processes, ranging from host defense to chronic inflammatory diseases.

Traditionally, monocytes were classified into three main subsets based on their surface expression of CD14 and CD16: classical (CD14^++^CD16^-^), intermediate (CD14^++^CD16^+^), and non-classical (CD14^+^CD16^++^) *(5)*. However, recent advances in single-cell technologies have unveiled a far more intricate monocyte landscape. Single-cell RNA sequencing (scRNA-seq) *(6, 7)* and mass cytometry (CyTOF) *(8)* have allowed researchers to appreciate the extensive transcriptional, phenotypic, and functional variability within these subsets. This diversity extends to the expression of pattern recognition receptors, chemokine receptors, and immune checkpoint molecules, which endow monocytes with specific functional capacities and responsiveness to pathogens and inflammatory stimuli. Classical monocytes are poised to engage in phagocytosis, acting as the first responders to invading pathogens, and have been implicated in various chronic diseases such as atherosclerosis *(9)*, cancer *(10)*, and rheumatoid arthritis *(11)*. Non-classical monocytes are more likely to be involved in patrolling and surveillance functions and are proposed to be protective against atherosclerosis *(12)* and cancer *(13)*. In contrast, data suggest that intermediate monocytes can be both proinflammatory or anti-inflammatory, suggesting vast functional heterogeneity *(14)*. The balance between these subsets, influenced by genetic and environmental factors, is critical for immune homeostasis and the promotion or inhibition of diseases.

Monocytes are crucial in the initiation and development of cardiovascular disease (CVD) *(15)*. Blood monocyte number is strongly associated with CVD in humans *(16)*, an association that has been reproduced in mouse models of atherosclerosis *(17–19)*. Traditional CVD risk factors, such as smoking and hyperlipidemia, can alter circulating monocyte numbers by increasing myelopoiesis *(20, 21)*, thereby promoting the progression of atherosclerosis. Consequently, the creation of a complete atlas of blood monocyte heterogeneity in healthy subjects and CVD risk states is crucial for developing targeted therapies in disease-promoting environments. As an initial step toward that end, we report here a comprehensive landscape of transcriptomes and cell surface phenotypes of over 400,000 circulating blood monocytes utilizing cellular indexing of transcriptomes and epitopes by sequencing (CITE-seq) from healthy individuals, smokers, and individuals with hyperlipidemia. Our analysis reveals perturbations associated with clinical and demographic variables that may promote the development of atherosclerosis and other chronic inflammatory diseases.

## Results

### High-dimensional CITE-seq improves the identification of monocyte subsets

In our pursuit to pinpoint unique protein and gene markers specific to monocytes, we conducted CITE-seq on monocytes obtained from 10 healthy volunteers, each prelabeled with a panel of 274 oligo-conjugate cell surface proteins (**Fig 1A**). This approach enables the comprehensive characterization of any cell type by simultaneously analyzing their transcriptome expression and cell surface protein expression, offering the potential for enhanced identification of distinct cell states *(22)*. Data from cells that met quality control and filtering criteria were integrated from a total of 62,692 cells across all volunteers, utilizing our previously published deep learning model *(23)* that had been adapted for the analysis of CITE-seq data. Subsequently, we employed clustering methods based on either RNA information alone or a combination of RNA and protein information, to assess whether this multimodal approach could enhance the resolution and identification of specific monocyte subgroups (**Fig 1B**). Based on RNA expression alone, UMAP visualization revealed 12 clusters, with 6 of these clusters being monocytes. However, when we applied clustering techniques to both RNA and protein modalities, we identified 14 clusters, of which 8 were monocytes. Notably, CITE-seq proved to be particularly effective in distinguishing non-monocyte populations from monocyte populations (**Suppl Fig 1A**).

**Figure 1.**
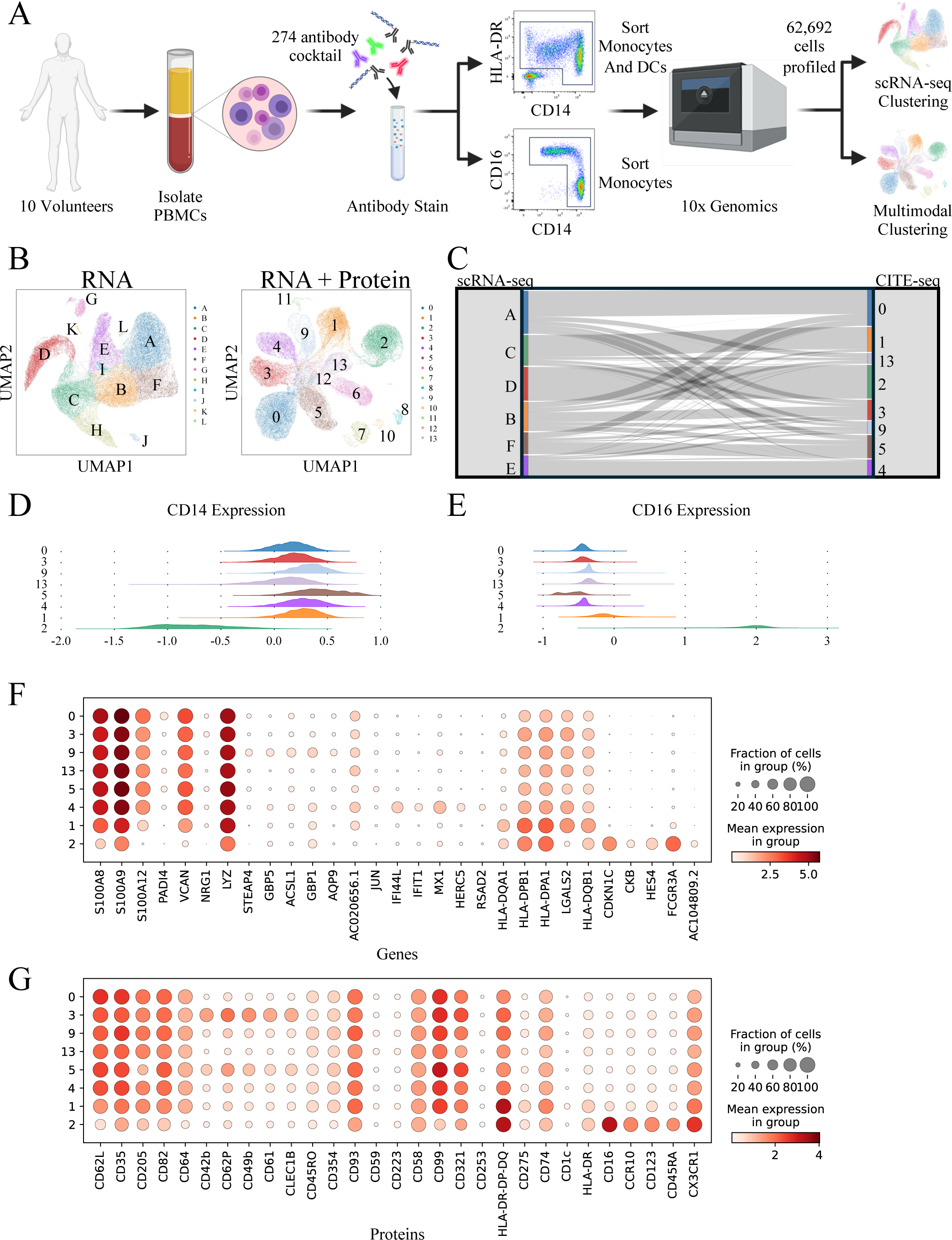
Multiomic analysis of human circulating monocytes identifies 8 monocyte subpopulations. (A) Overview of the experimental design. (B) UMAP visualizations using RNA (left) or RNA + protein (right) information. (C) Sankey plot mapping monocyte clusters identified in the RNA only analysis to the RNA + protein analysis. (D and E) Ridge plots showing the expression of the monocyte marker proteins CD14 (D) and CD16 (E). (F) Dotplot displaying the top genes for each monocyte subpopulation. (G) Dotplot displaying the top proteins for each monocyte subpopulation.

Further investigation, involving the mapping of RNA clusters to the CITE-seq clusters (**Fig 1C** and **Suppl Fig 1B**), revealed that several clusters previously identified based on RNA alone formed distinct clusters in the CITE-seq analysis. This strongly suggests that the inclusion of protein expression data substantially enhances the resolution and clustering of monocyte subsets. One of the most striking observations was the emergence of a monocyte population that displayed specific expression of numerous proteins, including CD62P, CD226, CD61, CD49b, and CD42b (**Suppl Fig 1C**). Interestingly, some populations were better defined by either RNA or protein expression (**Suppl Fig 1D**). Given the improvement in clustering and cell type identification through the combination of both modalities, it became evident that a multimodal analysis surpassed individual modality analyses. Therefore, we concluded that further characterization of monocyte subsets should encompass both modalities.

For the classification of each monocyte subpopulation into classical, intermediate, and nonclassical monocytes, we examined the expression levels of CD14 (**Fig 1D**) and CD16 (**Fig 1E**). Clusters 0, 1, 3, 4, 5, 9, and 13 displayed the highest CD14 expression, suggesting their classification as either classical or intermediate monocyte subpopulations. In contrast, cluster 2 exhibited minimal to no CD14 expression and distinctive CD16 expression, signifying it as the only nonclassical monocyte population within our dataset. Cluster 1 exhibited higher CD16 expression compared to other classical monocyte subsets, implying its representation as a CD14^++^CD16^+^ intermediate monocyte.

To maximize the potential of our multimodal analysis, we identified the top genes (**Fig 1F**) and proteins (**Fig 1G**) that defined each monocyte subpopulation. Clusters 0, 1, 3, 4, 5, 9, and 13 exhibited the classical monocyte genes *S100A8*, *S100A9*, and *S100A12* to varying degrees. These clusters also shared a similar protein expression profile, characterized by CD62L, CD35, and CD64. Cluster 3 (referred to hereafter as CD42b^+^) displayed a distinct protein expression profile, with specific expression of CD42b, CD62P, and CD49b. However, clusters 4 and 1 exhibited the most distinctive gene and protein profiles. Cluster 4 (referred to hereafter as IFN-responsive) displayed a unique gene expression profile, with specific expression of interferon-responsive genes such as *IFI44L* and *IFIT1*, suggesting involvement in antiviral activity. Cluster 1 (referred to hereafter as MHCII^hi^) was enriched in genes and proteins associated with the MHCII complex, including *HLA* genes and HLA-DR, DP, DQ proteins. Traditionally, these genes and proteins have been identified in dendritic cells (DCs) *(24)*. However, our analysis of the expression profiles of proteins CD275, HLA-DR, DP, DQ, and genes *HLA-DPB1* and *HLA-DQA1* revealed a noteworthy observation: cluster 6 (cDCs) also exhibits expression of these genes and proteins at levels similar to those in MHCII^hi^ monocytes. The distinction between these cell types was evident in the expression of CD14 and the DC-specific marker CD1c, highlighting that, despite their similarities, these cell types are indeed distinct (**Suppl Fig 1E**). The most distinct monocyte population was the nonclassical (cluster 2). As mentioned earlier, CD16 served as the most specific marker for this monocyte subset, along with several other proteins, including CCR10, CD123, and CD45RA.

### CITE-seq identifies new cell surface markers for monocyte subsets

Our comprehensive analysis unveiled distinctive cell surface protein markers that can contribute significantly to the classification, identification, and isolation of various monocyte subsets, as illustrated in **Figure 1G**. Our primary focus was to identify the most specific markers for both classical and nonclassical monocytes (**Fig 2A**). As anticipated, CD16, in agreement with existing literature, emerged as the most specific protein marker for distinguishing nonclassical monocytes. However, intriguingly, we found that CD99 surpassed CD14 in its effectiveness in distinguishing classical and intermediate monocytes from nonclassical monocytes. Additionally, CD275 and MHCII proved to be the optimal identifiers for the MHCII^hi^ monocyte population, while CD42b was the essential marker for the CD42b^+^ monocyte population. Nevertheless, for the IFN-responsive subset, the best classification was derived from RNA expression rather than unique protein profiles, therefore we could not identify cell surface proteins that can be used to characterize this subset. Consequently, specific cell surface proteins were integrated to construct a flow cytometry panel, aimed at effectively segregating select monocyte subpopulations (**Fig 2B**). Because previous studies have identified CD88 and CD89 as key markers for distinguishing between monocytes and dendritic cells *(24)*, we incorporated these proteins into our panel to ensure the purification of monocyte populations. Conventional flow cytometry validated the potential of combining CD99 and CD16 to notably enhance the separation of nonclassical monocytes from classical and intermediate populations. Furthermore, we were able to identify a distinct population of CD275^+^MHCII^+^ cells, corresponding to the MHCII^hi^ subset identified through our CITE-seq analysis. The remaining cells were constituted of predominantly classical monocyte subsets, where we pinpointed CD42b^+^ classical monocytes. However, other classical monocyte subpopulations remained challenging to delineate using conventional flow cytometry approaches. Subsequent back-gating of each population into the traditional CD14 and CD16 classification for monocytes revealed that CD16^+^CD99^-^ cells mapped back to nonclassical monocytes, MHCII^hi^CD275^+^ cells corresponded to intermediate monocytes, while both CD14^+^CD42b^+^ and the remaining CD14^+^ cells mapped back to subsets of classical monocytes (**Fig 2C**).

**Figure 2.**
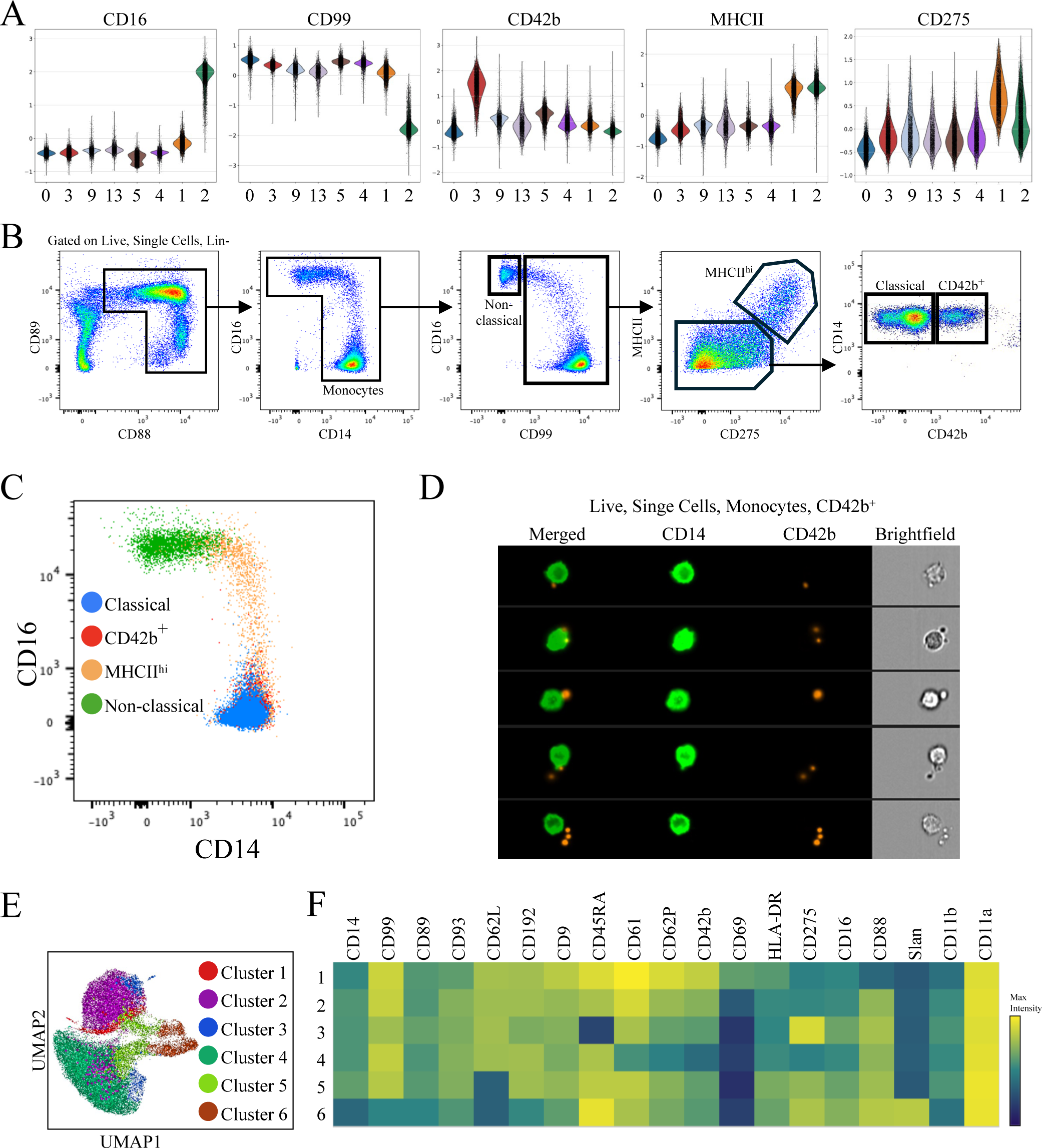
Identification and validation of monocyte subsets. (A) Violin plots depicting the expression of the proteins CD16, CD99, CD42b, MHCII, and CD275, which were the most discriminating proteins for each subpopulation identified in the CITE-seq analysis. (B) Optimal conventional flow cytometry gating strategy used to define and isolate non-classical, classical, MHCII^hi^, and CD42b^+^ monocyte subsets. (C) Mapping back of newly identified monocyte subsets to their historical classification based on CD14 and CD16 expression. (D) Imaging flow cytometry analysis used to identify CD42b^+^ cells as monocyte-platelet aggregates. (E) UMAP visualization of spectral flow cytometry data. (F) Heatmap showing the expression of select proteins across all clusters in the spectral flow analysis.

An intriguing observation arose during our analysis: many of the proteins expressed by the CD42b^+^ classical monocyte population are also found on platelets *(25)*. The possibility was raised, therefore, that this monocyte subset might represent monocyte-platelet aggregates, discernible only by analyzing protein expression through CITE-seq and not by transcriptome analysis alone. To address this possibility, we conducted imaging flow cytometry (**Fig 2D**). After gating on CD42b^+^ cells, we visually observed doublets consisting of platelets expressing CD42b and monocytes expressing CD14, thus confirming the presence of cells identified in CITE-seq that express CD42b and monocytes interacting with platelets.

For additional validation, we created a spectral flow cytometry panel based on top proteins identified in our CITE-seq analysis. When these cells were clustered based on the expression of these select cell surface proteins, six monocyte subsets were discerned (**Fig 2E**). The proteins defining these subsets were largely consistent with the populations we identified through CITE-seq, conventional flow cytometry, and imaging flow cytometry experiments. For instance, the non-classical population (cluster 6) was most distinct and exhibited very specific expression of CD16, CD88, CD45RA, and Slan. Similarly, classical subsets (clusters 1, 2, 3, and 4) were distinguished from nonclassical ones by their expression of CD99. Notably, we observed a classical subpopulation with specific expression of CD61, CD62P, CD42b, and CD69 (cluster 1), offering further independent validation of the existence of monocyte-platelet aggregates under healthy physiological settings. Clusters 3 and 5 displayed high expression of CD275 and MHCII, strongly suggesting that they correspond to the MHCII^hi^ population identified via CITE-seq.

In summary, our orthogonal approaches allowed us to identify and validate several biologically distinct monocyte subpopulations, with the most unique ones being the nonclassical, MHCII^hi^, IFN-responsive, and monocyte-platelet aggregates. Although we identified several classical subsets, they remain challenging to differentiate and isolate using existing technologies.

### Monocyte subpopulations exhibit unique pathways, trajectories, and transcriptional regulators

In our detailed analysis of monocyte subpopulations, we conducted various assessments to understand their unique pathways and developmental regulators. For classical monocytes (**Fig 3A**), as well as the non-classical, MHCII^hi^, and IFN-responsive monocyte subpopulations (**Fig 3B**), we analyzed differentially expressed (DE) genes. The predominant characteristic of classical monocytes was the upregulation of inflammatory and interleukin signaling pathways. The MHCII^hi^ subpopulation was notable for Th1 and Th2 signaling pathways, aligning with their MHCII complex-related genes and proteins crucial for T cell activation. Non-classical monocytes were associated with cell movement and fibrosis, while IFN-responsive monocytes exhibited distinct interferon signaling and antiviral response functions.

**Figure 3.**
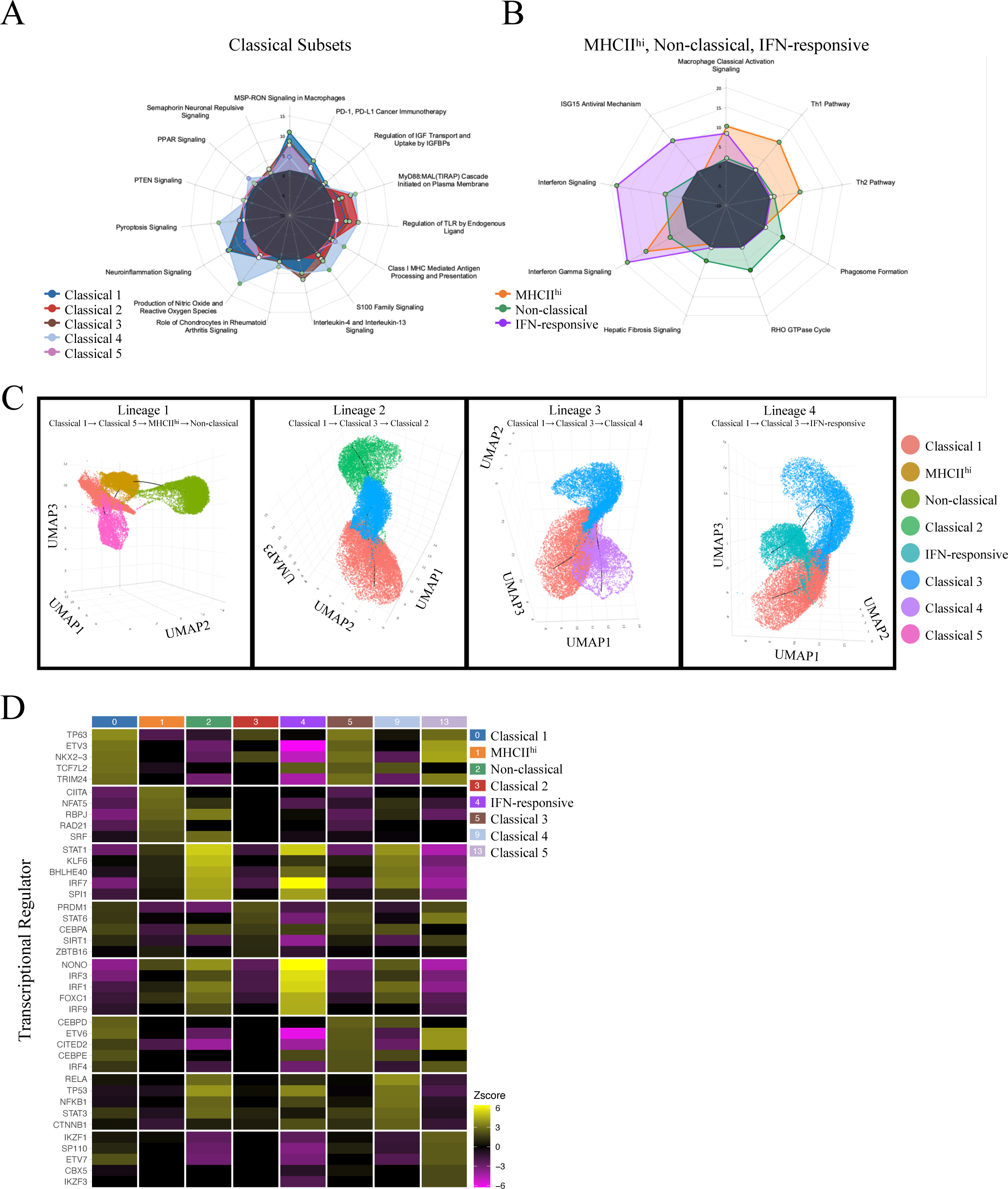
Monocyte subpopulations are functionally unique with distinct transcriptional regulation. Radar plots displaying top upregulated pathways determined by Ingenuity pathway analysis of classical (A), as well as MHCII^hi^, non-classical, and IFN-responsive (B) monocyte subpopulations. (C) Pseudotime trajectory analysis of monocyte subsets. (D) Heatmap showing top predicted transcriptional regulators for each monocyte subpopulation.

Previous studies indicate a sequential differentiation pathway for human blood monocytes, transitioning from classical to intermediate to non-classical subsets *(26)*. Building on this, we conducted a pseudotime analysis to delineate a more intricate differentiation trajectory among a more refined classification of blood monocytes (**Fig 3C**). The analysis uncovered four distinct developmental lineages. Each lineage begins with classical monocyte 1, which is likely to originate from the bone marrow. The first lineage follows the most observed progression, where classical monocytes differentiate into intermediate (termed MHCII^hi^ in our study) before ultimately terminally differentiating into non-classical monocytes. The second and third lineages pass through several classical monocyte subsets. However, the fourth lineage progresses through various classical subsets and culminates in the IFN-responsive subpopulation, a novel finding not previously reported.

To link the functionally distinct monocyte populations and their developmental pathways with potential transcriptional regulators governing these transitions, we conducted an upstream regulator analysis (**Fig 3D**). This revealed the following. All classical monocyte subsets shared a considerable overlap in transcriptional regulation, likely due to similarities in their gene expression profiles. In contrast, the MHCII^hi^, non-classical, and IFN-responsive subpopulations each exhibited distinct transcriptional regulators. Specifically, the MHCII^hi^ subpopulation was regulated predominantly by CIITA, a known master regulator of the class II major histocompatibility complex. In addition, the non-classical subpopulation showed inferred regulation by STAT1 and KLF6. Moreover, the IFN-responsive subpopulation appeared to be controlled by transcription factors associated with the innate immune response and antiviral activities, such as IRF3, IRF1, and IRF9, which are instrumental in regulating interferon expression.

### Identification of tissue-specific distributions of monocyte subpopulations by reference-based mapping

To identify and analyze the distribution of monocyte subpopulations across various human tissues, we leveraged the public dataset from Dominguez Conde et al *(27)*. This dataset consisted of blood, 10 non-lymphoid tissues, and 5 lymphoid tissues. We mapped all monocyte data to our blood monocyte CITE-seq atlas from **Figure 1**. However, we restricted our analysis to tissues containing over 100 monocytes, ensuring a sufficient cell count for assessing proportional distributions. Our final analysis was composed of cells from blood, bone marrow, liver, lung-draining lymph nodes, lung, skeletal muscle, and spleen (**Fig 4A**). Subsequent cell type prediction enabled us to accurately identify four monocyte subsets: MHCII^hi^, IFN-responsive, non-classical, and classical. Additionally, we observed a minor population of cDC2 (**Fig 4B**). The UMAP visualization of predicted cell types by tissue (**Fig 4C**) indicated the presence of all identified cell types across the studied tissues. We then quantified the proportion of each monocyte subpopulation in each tissue (**Fig 4D**), uncovering significant proportional differences linked to specific tissue reservoirs. Notably, skeletal muscle and liver displayed a higher proportion of non-classical monocytes, likely reflecting an abundance of terminally differentiated cells, while blood, bone marrow, and spleen predominantly harbored classical and possibly more immature monocytes. Intriguingly, the liver exhibited the highest proportion of MHCII^hi^ monocytes, and the skeletal muscle was the most enriched proportionally in IFN-responsive monocytes.

**Figure 4.**
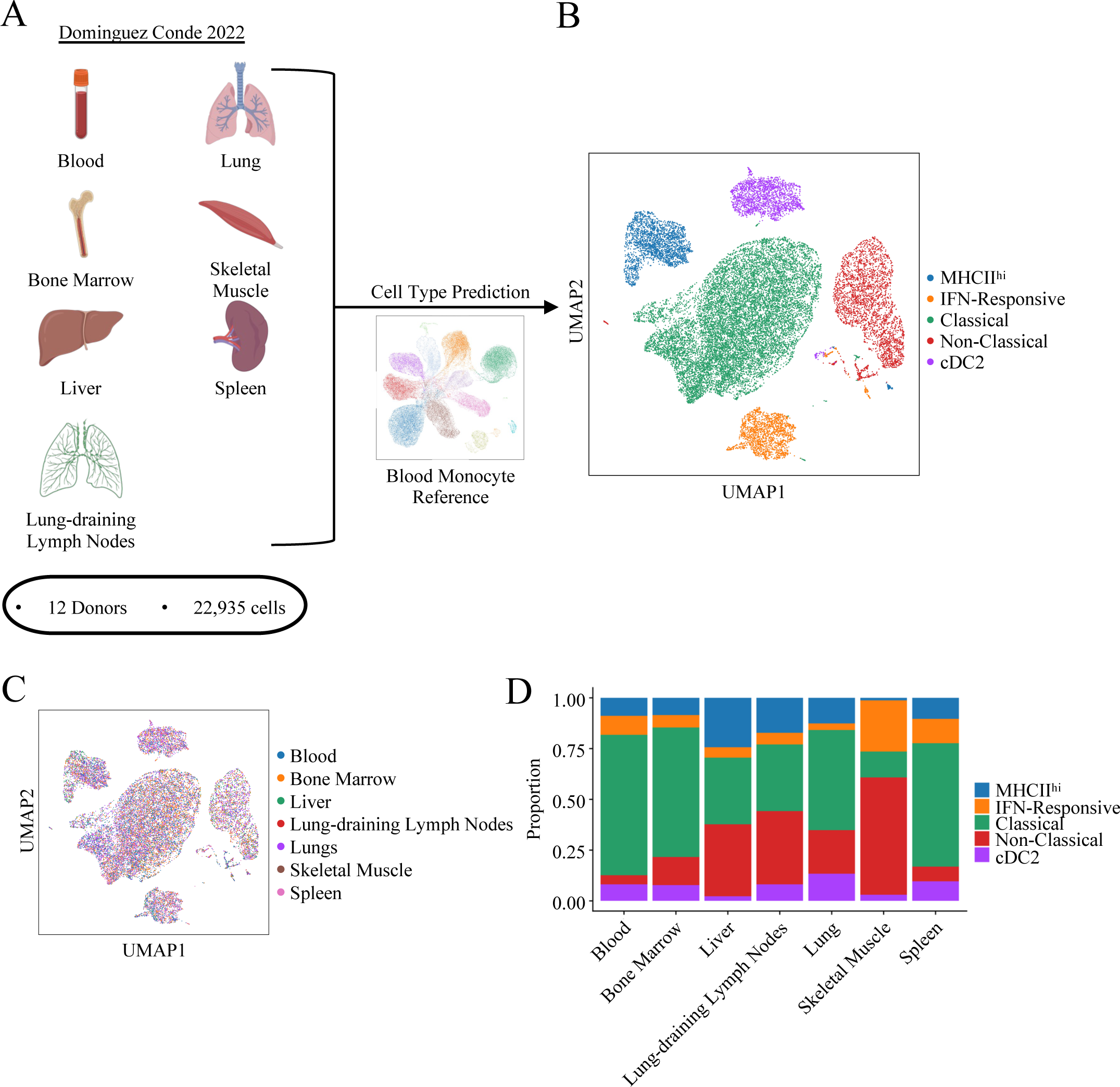
Tissue landscape of newly identified monocyte subpopulations. (A) Schematic illustrating data obtained and analyzed from Dominguez Conde (2022). (B) UMAP visualization of cell types identified in tissue data. (C) UMAP visualization colored by tissue origin. (D) Stacked bar graph displaying the proportional breakdown of each monocyte subset across all tissues included in the final analysis.

### Large-scale single-cell transcriptional profile reveals a comprehensive landscape of circulating monocytes

Our initial CITE-seq analysis unveiled significant phenotypic diversity within human circulating monocytes. To establish a comprehensive reference encompassing all monocyte subsets within a diverse population of humans, we expanded our scRNA-seq analysis to include 118 individuals, profiling a total of 351,499 cells (**Fig 5A**). While CITE-seq proved to be a superior method for profiling and identifying monocyte subsets, the cost and feasibility of applying this approach to all individuals was limiting. Consequently, we focused on transcriptional profiling of monocytes, delving deeper into their characterization.

**Figure 5.**
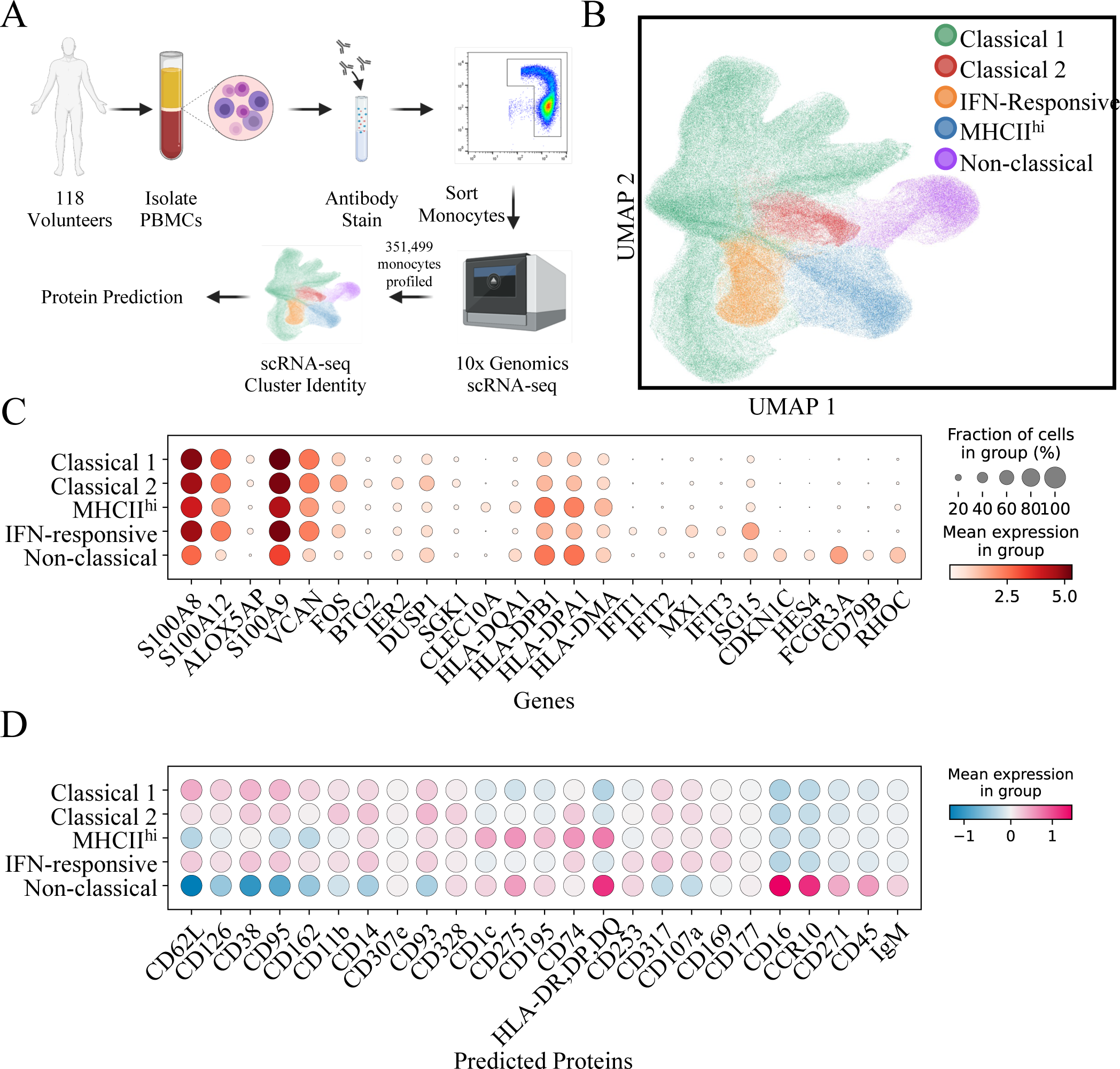
Comprehensive transcriptional landscape of circulating monocytes. (A) Experimental design for large scale profile of circulating monocyte subpopulation. (B) UMAP visualization of 351,499 cells profiled in this dataset. (C) Dotplot displaying top DE genes for each monocyte subpopulation. (D) Dotplot displaying top predicted proteins for each monocyte subpopulation.

In our analysis of this expanded dataset, we initially detected 12 cell clusters, with 9 of them representing monocytes (**Suppl Fig A**). Notably, our prior CITE-seq analysis had identified considerable similarities among classical monocyte subsets. As a result, we decided to combine classical monocyte clusters 0, 4, 6, 7, and 11, as the differential gene expression analysis revealed minimal distinctions between them (**Suppl Fig B and C**). This consolidation resulted in the identification of 5 major monocyte subpopulations. Additionally, we employed sciPENN, our recently published analytical method that uses a CITE-seq reference to predict and impute protein expression in scRNA-seq data *(28)*. Leveraging our CITE-seq analysis from **Figure 1**, we used these predictions to validate the consistency of phenotypic markers across the larger scRNA-seq dataset. Impressively, the protein predictions exhibited remarkable accuracy, reaffirming the expression of CD16 on nonclassical monocytes, CD14 and CD99 on classical monocytes, and MHCII and CD275 on MHCII^hi^ monocytes. However, CD42b was distributed across all populations, reflecting the challenge of identifying monocyte-platelet aggregates using scRNA-seq alone (**Suppl Fig D**).

Based on the top differentially expressed genes for each monocyte subpopulation from this analysis (**Fig 5C**), we observed a gene expression profile akin to our CITE-seq analysis in **Figure 1**. Classical 1 and 2 monocytes exhibited the highest expression of *S100A8*, *S100A9*, and *S100A12*, the MHCII^hi^ subpopulation expressed HLA genes, and the IFN-responsive population displayed increased expression of *IFIT1*, *IFIT2*, and *IFIT3*. Notably, the non-classical subpopulation exhibited specific expression of *FCGR3A*, the gene encoding CD16. Similarly, the inferred top differentially expressed protein markers for each population (**Fig 5D**) demonstrated consistency with our CITE-seq analysis.

In sum, our extensive scRNA-seq profiling of blood monocytes aligns cohesively with our initial CITE-seq cell type identification, reaffirming the robustness of our findings and the robustness and validity of predicting protein expression in large scRNA-seq datasets from reference CITE-seq profiling in a small subset.

### Demographic variables and clinical risk factors are associated with alteration in blood monocytes

To evaluate the impact of demographic factors and clinical risk variables, we conducted a Dirichlet regression analysis (**Fig 6**). Initially, we treated LDL-C as a continuous variable in our model to cover the full range of LDL-C levels (**Fig 6A**). This approach highlighted both pronounced and subtle variations. Notably, Black individuals were linked to a marked reduction in the MHCII^hi^ subpopulation (OR = 0.91; 95% CI = [0.83, 0.99]; P = 0.038) and a significant increase in the non-classical subpopulation (OR = 1.27; 95% CI = [1.14, 1.41]; P = < 0.001). Smoking showed a significant reduction in the non-classical subpopulation (OR = 0.82; 95% CI = [0.71, 0.94]; P = 0.004), with tendencies towards an increase in other subpopulations. Moreover, LDL-C levels were correlated with a non-significant increase in the non-classical subpopulation.

**Figure 6.**
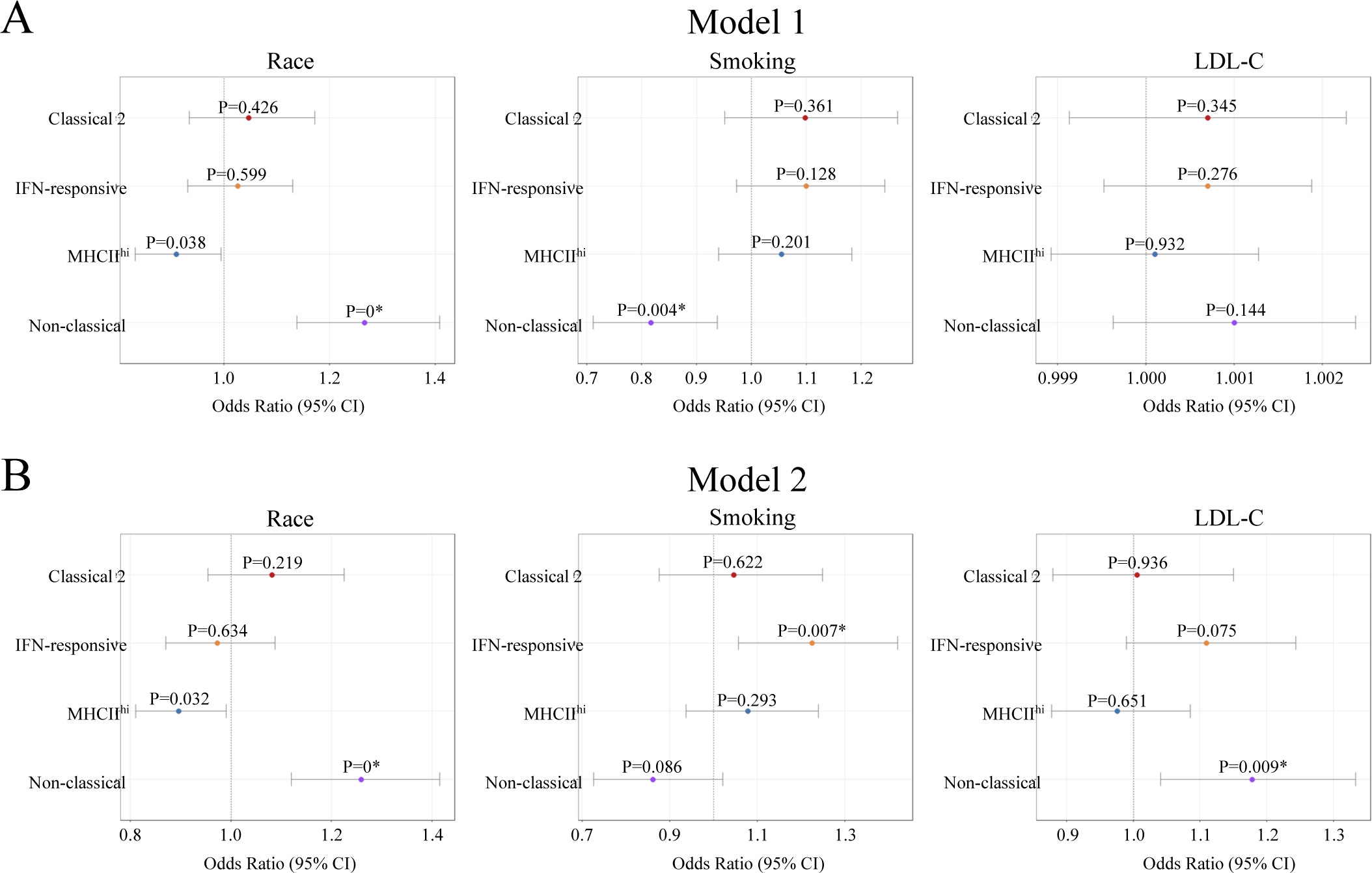
Association between demographic and clinical variables with monocyte distribution. Dirichlet regression analysis was performed to assess the relationship between age, sex, race, smoking, and LDL-C and alterations in the proportion of monocyte subpopulations. (A) Model 1 (n=91) used plasma LDL-C concentrations as a continuous variable. (B) Model 2 (n=74) used plasma LDL-C concentrations as a binary variable, where hyperlipidemic was considered LDL-C > 130mg/dL and LDL-C < 100mg/dL was considered normal.

In our second model (**Fig 6B**), we categorized participants based on LDL-C levels as a binary variable: high LDL-C (values above 130 mg/dL) and normal LDL-C (values below 100 mg/dL). This categorization helped in identifying changes associated with more extreme plasma LDL-C levels. Similar to the first model, the Black race was associated with an increase in the non-classical subpopulation (OR = 1.26; 95% CI = [1.12, 1.41]; P = < 0.001) and a decrease in the MHCII^hi^ subpopulation (OR = 0.90; 95% CI = [0.81, 0.99]; P = 0.032). Smoking continued to show a trend of a decrease in non-classical monocytes (OR = 0.86; 95% CI = [0.73, 1.02]; P = 0.086), but now a significant increase in the IFN-responsive subpopulation was evident (OR = 1.23; 95% CI = [1.06, 1.42]; P = 0.007). Notably, for high LDL-C levels, we observed significant increases in both the IFN-responsive and non-classical subpopulations (OR = 1.11; P = 0.075, OR = 1.18; P = 0.009). It should be noted that we observed no significant differences in monocyte frequency by age, sex, or HDL-C (**Table S1 and Table S2**).

### Similar monocyte subpopulations are present in mouse blood and alterations are associated with hyperlipidemia

We sought to strengthen the inferential basis of our findings of distinct human monocyte subpopulations and to extend our understanding of these subpopulations by investigating their presence, behavior, and responses to hyperlipidemic conditions in mouse models. We utilized both WT and LDL receptor-deficient (LDLr^-/-^) mice, feeding them either a standard chow diet or a Western diet for periods of 0, 8, or 16 weeks. At the end of these time points, we collected blood samples, sorted monocytes using FACS, and conducted scRNA-seq, analyzing a total of 31,523 cells (**Fig 7A**). Our approach involved reference-based mapping, using the human monocytes profiled with CITE-seq from **Figure 1** as the reference. This mapping allowed us to successfully identify counterparts in the mouse that were analogous to the human monocyte subpopulations (**Fig 7B**).

**Figure 7.**
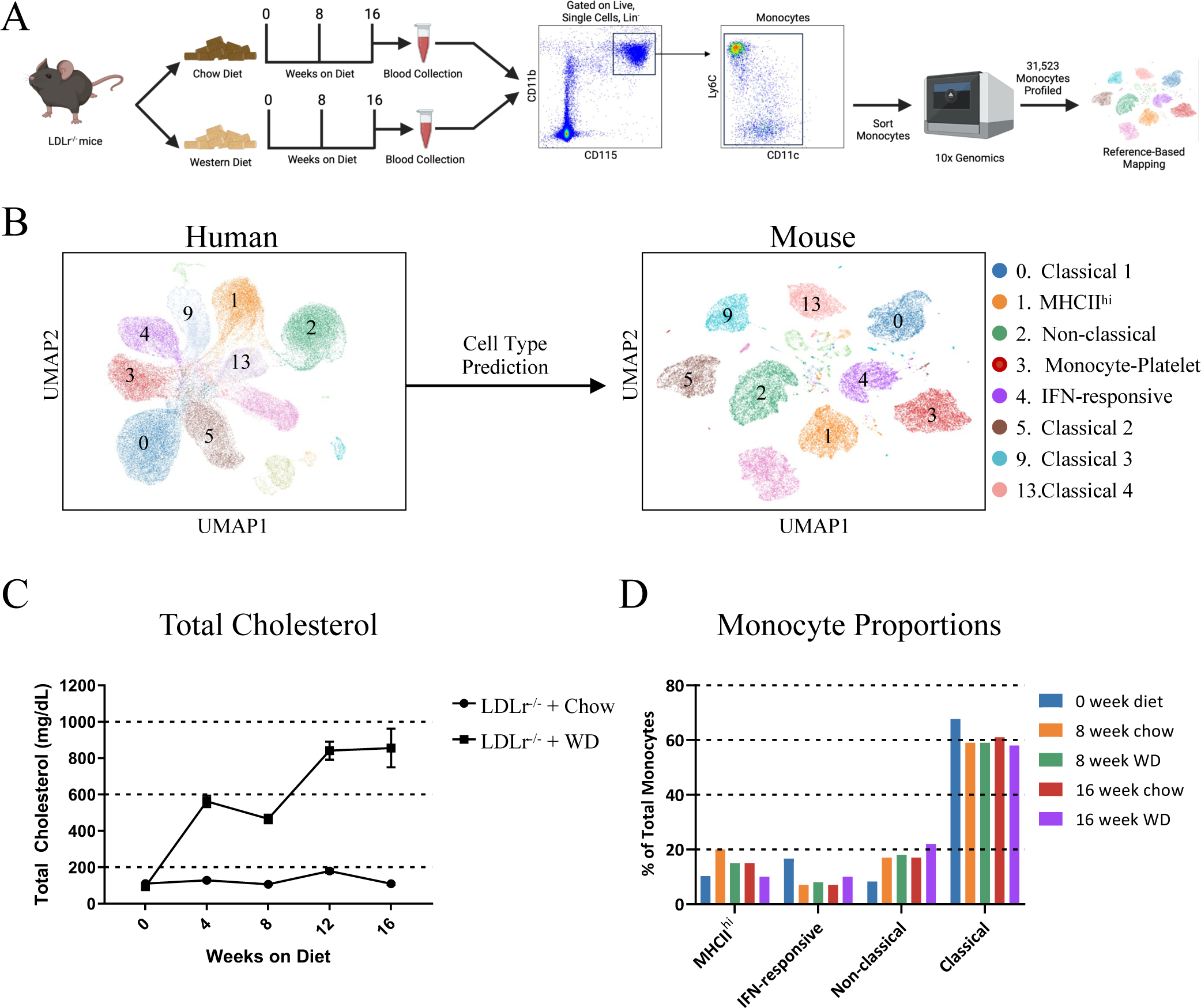
Single-cell RNA-seq profiling of mouse monocytes. (A) Experimental design for profiling mouse circulating monocytes in normal conditions or during hyperlipidemia. (B) Method for reference-based mapping used to identify human monocyte counterparts in mice. (C) Total cholesterol concentrations during diet time course. (D) Proportional breakdown of each monocyte subset in each diet condition.

We focused our final analysis on examining the behavior of monocytes in mice influenced by hyperlipidemic conditions. As expected, LDLr^-/-^mice on a Western diet had a significant increase in plasma cholesterol concentrations at 8 and 16 weeks (**Fig 7C**). We found a reduction in MHCII^hi^ monocytes in LDLr^-/-^ mice when compared to those fed a standard chow diet. Concurrently, there was a noticeable increase in both IFN-responsive and non-classical monocytes in these mice (**Fig 7D**). This pattern was very similar to the observations in human subjects, where we also found increased IFN-responsive and non-classical monocyte populations in humans with elevated LDL-C, as shown in **Figure 6**. These similarities in mice and humans underscore the utility of mouse models for in-depth exploration of monocyte functionality and their significance in both healthy and hyperlipidemic conditions. This comparative analysis not only validates the mouse model as a viable tool for studying monocytes in disease and their genetic regulation, but also helps in bridging our understanding of these cells across different species, particularly in the context of hyperlipidemia.

## Discussion

Here, we present the largest single multimodal dataset of circulating blood monocytes in humans, providing an in-depth reference of cellular heterogeneity and functional diversity within these cells. We made several novel discoveries, including the identification of transcriptionally and phenotypically distinct CD42b^+^, MHCII^hi^, IFN-responsive, and non-classical monocyte subsets. Additionally, we discovered several classical monocyte subsets. However, distinguishing and isolating these classical subsets proved challenging due to their overlapping gene and protein expression profiles, as well as similar inferred functionalities. Further expanding our analysis to include 118 individuals with CVD risk factors, we observed notable trends: an increase in non-classical monocytes linked to race, a reduction in non-classical monocytes associated with smoking, and a rise in both non-classical and IFN-responsive monocytes with hyperlipidemia. These findings suggest potential predispositions to CVD and other inflammatory diseases. Extending our research to hyperlipidemic mouse models, we noted consistent changes related to high cholesterol levels in mice, paralleling our human observations. This indicates that murine models can be instrumental in unraveling the disease-specific impacts of the monocyte subsets we have identified. Overall, this comprehensive dataset also serves as a crucial resource for the scientific community, facilitating the exploration of any perturbations linked to diseases involving monocytes.

Our work expands on recent efforts that have performed large-scale single-cell profiling and integration of myeloid cells across various tissues and disease states, utilizing technologies such as CyTOF *(8)*, high-dimensional flow cytometry *(24, 29)*, scRNA-seq *(6, 7)*, and more recently, CITE-seq *(30)*. These seminal studies have significantly advanced our understanding of myeloid cell diversity, prompting a reevaluation of the traditional CD14 and CD16 classification system for human monocytes. However, the generation of data across many studies and the integration of these data is prone to batch effect and computational artifacts. Although the findings from these studies largely agree on the substantial heterogeneity among monocytes, there is a lack of a clear consensus on the number of subsets present. That we have generated our data in the same lab using the same protocols limits this potential concern, providing a consensus atlas that is closer to the biological truth.

A major advantage of our multimodal profiling approach is its capacity to categorize monocyte subpopulations using a combination of gene and protein expression. This method yielded a more precise classification of each monocyte subset. A key finding was the identification of a CD42b^+^ classical monocyte subpopulation, distinguished by the expression of several platelet-associated proteins, including CD42b or platelet glycoprotein Ib alpha chain *(25)*. This discovery led us to investigate whether these were pure monocytes or monocyte-platelet aggregates. Our comprehensive flow cytometry analysis confirmed that incorporating protein data allowed us to detect a monocyte-platelet interaction, which is not discernible through transcriptional data alone. Notably, circulating monocyte-platelet aggregates are recognized as indicators of platelet activation, with an increased prevalence in patients with CVD *(31)*. Furthermore, platelets are known to facilitate the migration of monocytes into atherosclerotic plaques and promote a proinflammatory macrophage phenotype within these plaques *(32)*. These findings suggest that further research is essential to understand the implications of this interaction in health as well as in the pathophysiology of CVD. As noted, scRNA-seq alone is insufficient for identifying such interactions.

We also identified two other novel monocyte subsets, termed the MHCII^hi^ subpopulation and IFN-responsive. Our findings indicate that intermediate monocytes exhibit the highest expression levels of proteins and genes associated with antigen presentation, particularly those forming part of the MHCII complex *(33)*. Additionally, this subset is characterized by the specific expression of CD275, also known as ICOSLG. The counterpart of CD275, the costimulatory molecule CD278 or ICOS, is found on T cells *(34)*. This suggests that the MHCII^hi^ monocyte subpopulation plays a pivotal role in facilitating interactions with T cells, thereby promoting an adaptive immune response. The IFN-responsive subpopulation was characterized by having high expression of interferon-responsive genes that are involved in anti-viral activity. Interestingly, this population had no defining cell surface proteins and could only be defined by gene expression. It has been described that classical monocytes express the highest level of type 1 interferon genes *(35)*, and this signaling has been implicated in many immune diseases such as systemic lupus erythematosus.

Our analysis merging pseudotime with upstream regulatory predictions offers a compelling perspective on monocyte differentiation. Kinetic experiments suggest a sequential differentiation from classical monocytes, initially released from bone marrow, to intermediate, and ultimately non-classical monocytes *(26)*. This sequence, however, may oversimplify the differentiation process given the diverse monocyte subpopulations that have been identified. Our work reveals intricate developmental pathways, enriching the understanding of monocyte maturation beyond traditional models, and revealing four distinct developmental paths. While one pathway conforms to the well-documented progression from classical to intermediate to non-classical, two others transition through various classical monocyte subsets. Interestingly, a fourth pathway culminates in the differentiation into IFN-responsive monocytes, suggesting a more complex and nuanced understanding of monocyte maturation than previously appreciated. We also identified key transcriptional regulators that control monocyte differentiation. NR4A1 has emerged as a pivotal maser regulator, governing the shift from classical to non-classical monocytes and influencing their survival *(36, 37)*. Since we have shown that this is only one development trajectory, it is critical to start uncovering master regulators that control these newly identified monocyte lineages and how ablating specific subpopulations impacts inflammatory disease processes.

The critical involvement of monocytes in atherosclerosis development is well-established *(9)*. Several clinical studies have shown an association in monocyte number *(38, 39)* or shifts in the distribution *(40)* with CVD outcomes. However, these studies have used the CD14/CD16 classification system for monocytes, reducing the granularity of the findings. Moreover, the results have been inconsistent, with some studies linking higher classical monocyte counts to increased CVD risk and others suggesting associations with all monocyte subsets. Our extensive scRNA-seq dataset enabled us to identify specific changes in monocytes related to common CVD risk factors. Notably, smoking was linked to a reduction in non-classical monocytes, whereas hyperlipidemia was associated with an increase in non-classical and IFN-responsive monocytes. Although the pathophysiological significance of these alterations in relation to CVD remains to be fully elucidated, these alterations generate intriguing hypotheses for future research. Furthermore, the similarity in monocyte profiles between hyperlipidemic LDLr^-/-^ mice and humans suggests that in vivo studies in murine models will be informative to define the role of specific monocyte subpopulations in hyperlipidemia and its impact on atherosclerosis.

In conclusion, we provide a detailed characterization of human circulating monocytes in both normal and CVD risk conditions, uncovering phenotypically diverse monocyte subsets within the circulation and tissues. Notably, the prevalence of specific monocyte subsets changes in the context of CVD risk, suggesting promising targets for the prevention of CVD progression as well as valuable biomarkers of disease risk.

## Methods

### Human Studies

All human subjects research in this study, including the use of human tissues, conformed to the principles outlined in the Declaration of Helsinki. All patient information was de-identified. We enrolled 128 participants for this study that were recruited via direct advertising by the RecruitMe registry (https://recruit.cumc.columbia.edu/), which is supported by the Irving Institute for Clinical and Translational Research at Columbia University, and ResearchMatch.org, Craigslist, posting IRB-approved fliers on campus and community bulletin boards, newspaper advertising, and direct recruitment from the New York-Presbyterian Hospital Preventive Cardiology/Lipid clinic. We recruited healthy adults, individuals with type II diabetes, individuals with hyperlipidemia, and current and former smokers. All participants were generally healthy ranging from ages 29-73 years old and not on lipid-lowering medications. This study was approved by the CUIMC institutional Review board (IRB #AAAR5004). All study participants provided written informed consent. Inclusion criteria are adults able to consent. Exclusion criteria included active infection, known cardiometabolic disease, and other major comorbidities such as cancer.

### Plasma and Blood Collection

Blood was collected during the visit after a 12-hour fast. We submitted blood and serum samples to the Center for Advanced Laboratory Medicine (CALM) at Columbia University Medical Center for basic metabolic panel (assay 2921288), complete blood count with differential (assay 4646089), lipid panel (assay 4262402), cholesterol (assay 2921462), and Hemoglobin A1C (assay 2921552) analyses. Plasma LDL cholesterol levels were estimated using the Friedewald formula.

### PBMC Isolation

Blood from volunteers was collected in BD Vacutainer (sodium heparin) tubes (BD 367871) and immediately processed for peripheral blood mononuclear cell (PBMC) isolation. 4mL of blood was diluted to 5x with 16mL of DPBS + 2mM EDTA solution, for a final volume of 20mL, and carefully layered over 15mL of Ficoll-Paque Premium (GE Healthcare 17-5442-02) in 50mL conical tubes. Samples were centrifuged at 400 x g for 40 min at 20°C without brake. The top plasma layer was discarded and the PBMC layer was transferred to a new 50mL conical tube and subsequently washed with DPBS containing 2% heat-inactivated FBS, 5mM EDTA, 20mM HEPES, and 1mM sodium pyruvate (FACS buffer) by centrifugation at 500 x g for 10 mins at 4°C. PBMC pellet was then resuspended in 400uL of FACS buffer.

### Single-cell RNA-seq, CITE-seq, and Fluorescence-activated Cell Sorting

For CITE-seq, PBMCs were resuspended in 49uL of FACS buffer and 1uL of TruStain FcX (BioLegend: 422302) and blocked for 10 minutes at room temperature. Following incubation, lyophilized oligo-conjugated antibodies (**Table S2**) were reconstituted in 50uL of FACS buffer and were added to the cell suspension, and incubated for 30 minutes at 4°C. Then samples were washed 3 times in FACS buffer with centrifugation at 400g for 5 min at 4°C. PBMCs were incubated with a panel of fluorescently-conjugated antibody cocktail to enrich for monocytes, including CD14-AF488, CD16-PE-Cy7, HLA-DR-APC-eFluor 780, APC-labeled lineage markers (CD3, CD19, CD20, CD56, and CD66b) and DAPI. Monocytes were isolated by fluorescence-activated cell sorting (FACS) using a BD AriaII instrument by gating for live cells, then Lin^-^ and CD14^+^CD16^+^HLA-DR^+/-^ cells.

For scRNA-seq, PBMCs were incubated with a panel of fluorescently-conjugated antibody cocktail to enrich for monocytes, including CD14-AF488, CD16-PE-Cy7, HLA-DR-APC-eFluor 780, APC-labeled lineage markers (CD3, CD19, CD20, CD56, and CD66b) and DAPI. Monocytes were isolated by FACS using a BD Influx instrument by gating for live cells, then Lin^-^ and CD14^+^HLA-DR^+/-^ then CD14^+^CD16^+^ cells.

Cells were collected into a 15mL conical tube containing 7mL of RPMI media + 10% FBS and the final cell suspension was immediately subjected to single-cell encapsulation. All single-cell experiments were performed at the Single Cell Core Facility at the JP Sulzberger Columbia Genome Center using the Chromium Single Cell Gene Expression system (10x Genomics). Samples were prepared following 10x Genomics 3’ v3 protocol according to the manufacturer’s instructions for cDNA amplification. 5000 cells and 100M reads were targeted per sample.

### CITE-seq Library Preparation

CITE-seq libraries were prepared as previously described *(22, 41)* following 10x Genomics 3’ v3 protocol according to the manufacturer’s instructions for cDNA amplification using 0.2 μM of ADT additive primer (5’CCTTGGCACCCGAGAATTCC). The supernatant from the 0.6x SPRI cleanup was saved and purified with two rounds of 2x SPRI, and the final product was used as a template to produce ADT libraries. Antibody tag libraries were generated by PCR using Kapa Hifi Master Mix (Kapa Biosciences KK2601), 10 mM 10x Genomics SI-PCR primer (5’AATGATACGGCGACCACCGAGATCTACACTCTTTCCCTACACGACGCTC), and Small RNA RPIx primer (5’CAAGCAGAAGACGGCATACGAGATxxxxxxGTGACTGGAGTTCCTTGGCACCCGAG AATTCCA with xxxxxx denoting one of the four following sequences: CGTGAT, ACATCG, GCCTAA, TGGTCA). Following amplification, Antibody tag libraries were cleaned up with 1.6x SPRI. Subsequently, ADT quality was verified using a DNA high sensitivity assay on an Agilent 2100 bioanalyzer.

### ScRNA-seq and CITE-seq data processing

Fastq files were processed by Cell Ranger 5.0.1 from 10X genomics to generate count matrices of unique molecular identifiers (UMIs). Reads were mapped to the human reference genome GRCh38 and transcriptome annotated by GENCODE v32 from 10X prebuilt reference package version 2020-A. For CITE-seq libraries, an additional feature barcode reference of 283 TotalSeq-A antibody-derived tags was included. As some gene IDs from the reference share the same gene symbol, a suffix number was added to the gene symbol for each duplicate gene ID.

RNA level and cell level filtering were performed on each sample. RNAs were included if they were expressed in ≥10 cells. Cells were included if they had ≥200 RNAs expressed. As monocytes are a relatively homogeneous population with tight distributions of RNA and UMI counts, we applied stringent cutoffs on the maximum number of RNAs and the maximum number of UMIs allowed for each sample. We first calculated the 75^th^ percentile RNA count for each sample, and set the cutoffs to 1) 3,000 RNAs and 20,000 UMIs if the 75^th^ percentile RNA count ≥2,400, 2) 2,500 RNAs and 20,000 UMIs if the 75^th^ percentile RNA count ≥1,800 but <2,400, and 3) 2,000 RNAs and 15,000 UMIs otherwise. In addition, cells with >10% of reads mapped to mitochondrial genes were excluded from downstream analysis.

Filtered UMI count matrices from 118 human monocyte scRNA-seq datasets were used as input to CarDEC *(23)*, a deep learning-based tool for batch effect correction, gene expression denoising, and cell clustering. Sample ID was set as the batch variable, and 2,500 highly variable RNAs were selected for clustering analysis. A total of 12 clusters were identified, including 9 monocyte clusters, T cells, B cells, and platelets. Monocytes were retained for downstream analysis and were collapsed into 5 subpopulations based on transcriptome similarity.

Cells were visualized by Uniform Manifold Approximation and Projection (UMAP). Briefly, “tl.pca” function from Scanpy package *(42)* was applied to low-dimensional embedding from CarDEC, followed by “pp.neighbors” and “tl.umap” functions.

### Filtering and CarDEC_CITE clustering of CITE-seq data

Feature level and cell level filtering were performed on each sample. RNAs and proteins were included if they were expressed in ≥10 cells. Cells were included if they had ≥200 RNAs expressed. The maximum number of RNAs and the maximum number of UMIs from RNA allowed varies slightly across samples. In addition, cells were filtered if there were >50 UMIs from control antibodies. A maximum of 10% reads were allowed that were mapped to mitochondrial genes.

CarDEC_CITE, a modified version of CarDEC was applied to filtered RNA and protein UMI count matrices from 15 human monocyte CITE-seq samples. Sample ID was used as the batch variable. 2,000 highly variable RNAs and 100 highly variable proteins were selected for clustering analysis. A total of 14 clusters were identified, including 8 monocyte clusters, NK cells, dendritic cells, mast cells, endothelial cells, and platelets. Monocyte clusters were further collapsed into 4 subpopulations based on transcriptome similarity.

To assess whether multimodal data improves resolution of monocyte subclusters, we applied CarDEC on RNA subset of the CITE-seq data using 2,000 highly variable RNAs. A total of 6 monocyte clusters were identified along with non-monocyte populations. Cells were visualized by UMAP as described in scRNA-seq analysis.

### SciPENN protein prediction

We utilized sciPENN *(28)*, a deep learning tool, to predict protein expression in the larger scRNA-seq dataset leveraging information learned from CITE-seq data. Specifically, RNA and protein expression from CITE-seq were used as the training set, and RNA expression from scRNA-seq was used as the test set in sciPENN. Protein expression in scRNA-seq data was predicted in a z-score scale. Differential expression of predicted protein expression among monocyte populations was performed using Wilcoxon rank-sum test in “tl.rank_genes_groups” function from Scanpy package.

### Differential expression analysis in scRNA-seq and CITE-seq data

Raw RNA counts were normalized by “pp.normalize_total” function in Scanpy package with total count set to 10,000 and then log-transformed with pseudo-count of 1. Raw protein counts were normalized using centered log ratio method implemented in “prot.pp.clr” function from MUON package *(43)*. Differentially expressed RNAs and proteins were identified using the Wilcoxon rank-sum test from “tl.rank_genes_groups” function in Scanpy package. RNAs and proteins were required to be expressed in ≥25% of cells from either group. P-values were adjusted by the Benjamini-Hochberg method. Cell types were annotated using the top differentially expressed RNAs and proteins with adjusted P-value <0.05.

### Pseudotime analysis of human monocytes

Low-dimensional embeddings from CITE-seq data were utilized to infer pseudotime trajectories of human monocytes. UMAP was generated using the same process as in scRNA-seq analysis and setting the number of components to 5. The 5-dimensional UMAP coordinates for monocytes were used as input to Slingshot *(44)*, a program for trajectory inference. “Classical monocyte 1” was used as the starting cluster. Inferred trajectories were visualized in the first 3 UMAP dimensions using plotly R package.

### Reference-based mapping of human monocytes from public scRNA-seq data

Raw count matrix was downloaded from Cross-tissue Immune Cell Atlas *(27)* using the myeloid compartment data. Monocyte expression was extracted by selecting “Nonclassical monocytes” and “Classical monocytes” from the “Manually_curated_celltype” annotation. ItClust *(45)*, a transfer learning method for cell type classification, was used to predict monocyte subpopulations in the public data. RNA expression from human monocyte CITE-seq data was used as reference along with cell type labels including MHCII^hi^, IFN-responsive, classical, and non-classical monocytes and cDC2. Predicted clusters with a confidence score above 80% were included in downstream analysis. Proportions of predicted cell types were calculated in tissues with over 100 monocytes.

### Pathway Analysis

Cell-type specific differentially expressed (DE) genes, with a Benjamini-Hochberg adjusted p-value of less than 0.05 and log_2_ fold change greater than 0.58 and less than -0.58, were uploaded into Ingenuity Pathway Analysis (IPA) software (QIAGEN) *(46)*. IPA analysis reported the -log_10_ p values and Z-scores of canonical pathways and upstream regulators. The highest predicted upregulated canonical pathways and upstream regulators with a -log_10_ p value > 1.3 (p-value < 0.05) for each cell population were selected for the respective analysis and visualized with either a heat map or radar plot.

### Human regression analysis

Human monocyte scRNA-seq datasets from 116 subjects were first merged with their demographic information and clinical outcomes. The current analysis included 99 White or Black individuals with a glucose level equal to or less than 150 mg/dL and an HDL-C level equal to or less than 150 mg/dL. For more precise smoking measurement, cotinine levels were also taken into account to identify whether smoking use is consistent with cotinine levels. Smokers with a cotinine level greater than or equal to 50 ng/mL and non-smokers with a cotinine level less than or equal to 500 ng/mL were considered. To further investigate the effect of the two groups with normal or high level of LDL-C, individuals with LDL-C levels below 100 mg/dL were classified as the normal LDL-C group, and those with levels above 130 mg/dL as the high LDL-C group. Clinical and demographic baseline characteristics are provided in **Supplemental Table 5**. Dirichlet regression models were then utilized to identify the proportional composition of monocytes between groups, using the DirichletReg 0.7–1 package *(47)* in R version 4.1.0.

Fractional monocyte sub-populations were calculated for individuals based on five pre-classified cell types: Classical 1 monocytes, Classical 2 monocytes, MHCII^hi^ monocytes, IFN-responsive monocytes, and non-classical monocytes. In this analysis, Classical 1 monocytes were used as a baseline for alternative parameterization, making Dirichlet regression interpretable in a manner similar to multinomial logistic regression. Conceptually, the traditional Dirichlet regression follows the probability density function: 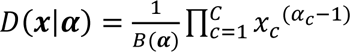, where *B* denotes the multinomial beta function, while ***x*** represents the proportions of cell types, and ***α*** signifies the model parameters*(48)*. Both ***x*** and ***α*** have a length equivalent to C, which is 5 for this analysis. In the alternative parameterization, the expected values of the components are represented as a vector μ, with each element μ_*c*_ ranging from 0 to 1, and the sum of μ equals to 1, and φ is a precision parameter. The group means and the precision parameter are explicitly modelled by defining *α*_0_ = φ and *α*_*c*_ = μ_*c*_*φ*. This leads to regression models where μ modeled based on the multinomial logit function, and φ is modeled using the log-link function. Leveraging the features of the alternative parameterization, the comprehensive model was initially adjusted for age, sex, race, smoking use, HDL-C, and LDL-C. It was then fitted to both the mean and precision models, using the continuous LDL-C variable from all subjects. The LDL-C variable was then replaced with a binary LDL-C variable to identify differences in effects between the normal and high LDL-C groups. This fully adjusted model was also used to estimate both the expected values and their precision.

### Imaging Flow Cytometry

Fresh PBMCs were isolated as described above. Subsequently, cells were blocked for 10 minutes at room temperature in TruStain FcX. Cells were incubated with the following antibodies for 20 mins at 4°C: CD14-FITC, CD16-PE-Cy7, HLA-DR-APCeFluor 780, CD42b-BV510, Lineage-APC. For imaging flow cytometry, cells were acquired on the ImageStream^X^ MkII imaging cytometer. The signal was measured in the following channel: Ch1 brightfield camera 1, Ch 2 488-560nm, Ch 560-595nm, Ch 6 745-785, Ch7 435-505, Ch 8 505/570, Ch 9 brightfield, Ch 11 642/745, Ch 12 745/785. The following gating logic was used: single cells were selected using area brightfield against aspect ratio brightfield, in focus cells were selected by using the gradient RMS feature, live cells were gated on the negative in Ch 7 (DAPI). Lineage-cells were selected by selecting the negative population in Ch 11 (APC), monocytes were gated by Ch 12 (APC-eflour 780) vs Ch 2 (FITC). Monocytes were purified by gating on CD14 (FITC) and CD16 (PE-Cy7). CD42b+ were selected in Ch 8 (BV510). From that population, images were examined to assess the presence of monocyte-platelet aggregates at 40x magnification. Data was analyzed on IDEAS v6.2.

### Conventional and Spectral Flow Cytometry

Fresh PBMCs were isolated as described above. Subsequently, cells were blocked for 10 minutes at room temperature in TruStain FcX. Cells were then incubated with appropriate antibodies, depending on the analysis being performed, and brilliant stain buffer (BD Biosciences) for 30 minutes at 4°C, then washed three times. For conventional flow cytometry cells were analyzed on BD FACSAria II and for spectral analysis cells were analyzed on a 5L Cytek Aurora 5 laser spectral analyzer. Fcs files for conventional and spectral flow cytometry were exported and analyzed using FlowJo v10.8.1.

### Mouse Experiments

All animal experiments were approved by the Institutional Animal Care and Use Committee (IACUC) of Columbia University (protocol AC-AABQ5576). C57BL/6J (Strain No: 000664) and LDLr^-/-^ (Strain No: 002207) were obtained from Jackson Lab. At 8 weeks of age, mice were either sacrificed or put on a Western diet (Inotiv TD.88137) or chow for 8 or 16 weeks.

### Single-cell Preparation of Mouse Monocytes

At the termination of the diet period, mice were anesthetized with ketamine/xylazine, and blood was collected by cardiac puncture. PBMCs were isolated by lysing red blood cells with RBC lysis buffer (BioLegend: 420302) following the manufactured protocol. Cells were then blocked with TruStain FcX PLUS (BioLegend: 156604). Subsequently, cells were stained with the following cocktail of antibodies: Lineage-APC, CD11b-PE-Cy7, CD11c-BB515, CD115-PE, and Ly6C-BV605, and washed three times. Monocytes were isolated by FACS using a BD FACSAriaII instrument by gating for live cells, then lineage negative, followed by CD115+ and CD11b+, and monocytes were identified as being Ly6C+/-. All scRNA-seq experiments were performed at the Single Cell Core Facility at the JP Sulzberger Columbia Genome Center using the Chromium Single Cell Gene Expression system (10x Genomics). Samples were prepared following 10x Genomics 3’ v3 protocol according to the manufacturer’s instructions for cDNA amplification. 5000 cells and 100M reads were targeted per sample.

### Mouse cell type prediction analysis

The three blood monocyte mouse scRNA-seq datasets and the five Ldlr^-/-^ mouse scRNA-seq datasets were first processed with Seurat version 4.3.0 *(49)* in R software version 4.1.0. To control the quality of the data, genes expressed in fewer than ten cells, as well as cells expressing fewer than two hundred genes and containing more than ten percent of mitochondrial genes, were discarded. Each single cell dataset underwent further filtration based on sequencing depth, with the comprehensive parameters outlined in **Supplemental Table 6**. The eight pre-processed datasets were then merged, and biomaRt version 2.48.3 *(50)* was used to map the mouse genes to their corresponding human orthologues, excluding any genes that lacked orthologues. ItClust version 1.2.0*(45)* was utilized with Python version 3.6.8 to predict mouse cell types. The human CITE-seq data with pre-labeled cell types was used as a reference dataset to predict mouse cell types and identify similarities in monocyte subpopulations across different species. Using ItClust, 1,770 highly variable genes from the mouse dataset were selected, and the corresponding gene expression matrix from the human data was used to cluster the mouse cells and predict their cell types. Fifty principal components and fifteen nearest neighbors were used to generate Uniform Manifold Approximation and Projection (UMAP) for the mouse data using Scanpy version 1.8.1*(42)*. For the differential gene expression analysis, mouse gene expression counts were normalized by the total counts, multiplied by 10,000, and then natural log transformed after adding a pseudo-count of 1. The Wilcoxon Rank Sum test was used to identify the top differentially expressed genes obtained by comparing a cluster to all of the other clusters. Mouse genes expressed with at least a 1.5-fold difference and a Benjamini-Hochberg-corrected p-value of less than 0.05 were considered as differentially expressed for each predicted cell type.

### Plasma Cholesterol Measurement

Plasma was collected by retroorbital bleed at indicated time points. Total plasma cholesterol concentrations were determined using the Infinity Cholesterol Reagent (Thermo Fisher Scientific: TR13421) following the manufacturer’s recommendations.

## Supplementary Materials

Fig S1 and S2

Table S1 to S6

## Acknowledgments

The scRNA-seq and CITE-seq (10x Genomics) were performed in the JP Sulzberger Columbia Genome Center, supported in part through the National Institutes of Health/National Cancer Institute Cancer Center Support Grant P30CA013696, and used the Genomics and High Throughput Screening Shared Resource. Human CITE-seq, conventional flow cytometry, imaging flow cytometry, and mouse scRNA-seq flow cytometry experiments described in this article were performed in the Columbia Stem Cell Initiative Flow Cytometry core facility at Columbia University Irving Medical Center. FACS sorting for scRNA-seq and spectral flow cytometry analysis and FACS sorting were performed in the Flow Cytometry Core at Columbia Center for Translational Immunology, supported in part by the NIH award S10OD020056.

## Funding

National Institute of Health grant R01HL113147

National Institute of Health grant R01HL150359

National Institute of Health grant R01HL166916

National Institute of Health grant R01HL169766

National Institute of Health grant 2T32HL007343

National Institute of Health grant R01HL151611

National Institute of Health grant R01HL168174

National Institute of Health grant K99HL153939

Irving Scholar Award from Columbia University Irving Institute for Clinical and Translational Research

American Heart Association grant 23PRE1026409

American Heart Association grant 19POST34450233

## Author contributions

Conceptualization: ACB, HP, HZ, ML, MPR

Methodology: ACB, CX, EK, HY, HP, MK, HZ, ML, MPR

Investigation: ACB, CX, EK, HZ, HZ

Visualization: ACB, CX, EK, LYZ, MK

Funding acquisition: ML, MPR

Project administration: LSR, ML, MPR

Supervision: ML, MPR

Writing – original draft: ACB, CX, EK, MPR

Writing – review & editing: ACB, CX, EK, HZ, ML, MPR

## Competing interests

None

## Data and Materials Availability

All data will be released on the Gene Expression Omnibus (GEO) upon acceptance of this article.

## Notes

### Competing Interest Statement

The authors have declared no competing interest.

